# FlowWeb: a free, web-based platform for flow cytometry data analysis

**DOI:** 10.64898/2026.04.16.717288

**Authors:** Menno ter Huurne, Rustem Salmenov, Amit Mandoli

## Abstract

Flow cytometry is widely used for high-throughput single-cell analysis. However, its data analysis relies on either costly commercial software or programming-intensive open-source tools. To bridge this gap, we developed FlowWeb, a freely accessible, web-based platform that combines the flexibility of the R/Bioconductor ecosystem with an intuitive graphical user interface. FlowWeb enables integrated workflows for data handling, quality control, gating, visualization and statistical analysis within a unified environment.

FlowWeb integrates raw data, metadata, and analytical state within synchronized Bioconductor structures, enabling coherent analysis and visualization workflows. FlowWeb supports both manual and automated data-driven gating workflows. To evaluate its performance, we applied FlowWeb to an in-house flow cytometry dataset and compared its automated cell cycle and gating workflows to established commercial tools. FlowWeb’s automated cell cycle workflow produced consistent and reproducible results across replicates and demonstrated high concordance with reference analyses, highlighting the platform’s robustness. FlowWeb’s advanced visualization tools include a wide range of fully customizable individual, overlay, and statistical plots. To enhance usability and reproducibility, the FlowWeb platform provides optional user-accounts that allow storage of reusable configurations, including quality control presets, gating definitions, and plot templates.

By lowering technical barriers without compromising analytical rigor, FlowWeb facilitates accessible, reproducible, and scalable flow cytometry data analysis for a broad range of users in research and clinical settings.

## Introduction

Flow cytometry is a cornerstone technology in modern biomedical research and clinical diagnostics, enabling rapid, high-throughput measurement of cellular phenotypes at the single-cell level. It is used worldwide across diverse fields including immunology, oncology, and hematology, where precise characterization of complex cell populations is essential. The analysis of flow cytometry data is a critical step in the workflow, often requiring specialized expertise and dedicated software environments.

Several commercial software packages, such as FlowJo (BD Biosciences), Kaluza (Beckman Coulter), and FCS Express (De Novo Software), provide comprehensive solutions for data visualization, gating, and statistical analysis. While these platforms are powerful and widely used, they are typically associated with substantial licensing costs and may impose limitations on accessibility, reproducibility, or integration with modern data science workflows. In parallel, a rich ecosystem of open-source, R-based tools (e.g., flowCore, flowWorkspace, openCyto) provides highly flexible and reproducible alternatives. However, effective use of these packages generally requires programming expertise, which remains a barrier for many experimental researchers who rely on graphical user interfaces.

To address these challenges, we developed FlowWeb, an interactive, web-based platform for flow cytometry data analysis that combines the flexibility of the R/Bioconductor ecosystem with an intuitive graphical user interface. FlowWeb enables reproducible, scalable, and user-friendly analysis by integrating data handling, visualization, gating, and statistical evaluation within a unified environment. By leveraging open-source infrastructure and a modular architecture, it lowers the barrier to advanced cytometric analysis while maintaining analytical rigor and transparency.

## Data handling/architecture

FlowWeb is built on a layered data model that separates cytometry data, metadata, and analytical state into distinct but synchronized components.

Uploaded FCS files are represented as *flowFrame* objects using the Bioconductor flowCore infrastructure [1]. Each file is assigned both a stable internal identifier and a user-friendly label, allowing reliable data handling while maintaining clarity in the interface.

Initially, uploaded FCS-files are grouped in flowSets based on their (shared) detector configurations, which facilitates data-preprocessing (such as Transformations). For each *flowSet* a corresponding *GatingSet* is created. The *flowSet* stores the data and derived populations, while the GatingSet [2] captures the hierarchical gating structure and ensures that analytical steps are applied consistently across samples. The software keeps the GatingSet registry automatically synchronized with the flowSet registry, including creation of new GatingSets for new flowSets and removal of obsolete GatingSets when flowSets are deleted. This design facilitates the processing of multiple files with distinct antibody panels and/or detector configurations in one session, reflecting the practical organization of flow cytometry studies, where multiple staining panels are commonly analyzed in parallel.

Sample annotation data can be imported from spreadsheets and are integrated with cytometry data in a central interactive table. This table links samples, metadata, and derived populations, enabling intuitive navigation and analysis. The application is implemented in Shiny, which manages user interactions and keeps all components synchronized.

Importantly, uploaded cytometry data and sample annotation files are processed within the active session and are not persistently stored on the server. This session-based handling minimizes long-term data retention while maintaining interactive performance.

## The FlowWeb Interface

FlowWeb is built as an interactive shiny environment embedded within a website, combining a reactive analysis interface with the accessibility of a web-based platform. All computations are performed on a secure, EU-based server, ensuring centralized processing and facilitating compliance with data protection standards. In addition to the core analytical workspace, the website provides tutorial material to guide users through the main features of the software, with further tutorials expected as the platform develops. The site also includes a personalized user environment, “My FlowWeb,” which enables storage and reuse of analysis-related settings and will be discussed in more detail at the end of the manuscript. FlowWeb is hosted as a web platform and is anticipated to expand over time, with additional functionality, interface improvements, and supporting resources.

### The main table

#### Architecture and Interaction

Uploaded FCS-files are integrated into a reactive table that serves as the main interface for the dataset. Root rows correspond to imported FCS files, whereas child rows represent gated subpopulations derived from a parent sample. Under the hood, derived populations store their immediate parent in their metadata, enabling reconstruction of the full lineage tree. This parent-child structure is used to calculate table indentation, build collapsible hierarchies in the interface, propagate metadata, and support cascaded editing or deletion of (sub-) populations. In addition, parent-child structure also serves as the substrate for summary statistics. Event counts are taken directly from each flowFrame, and the number of events as percentages of its parent (%Par) and its root sample (%Tot) are computed from the parent-child hierarchy.

The table is highly interactive and supports multiple levels of navigation, selection, and data exploration. Hierarchical relationships between samples and derived populations can be expanded or collapsed, while context menus provide access to advanced operations such as file inspection, gate application, and data export. Users can directly visualize and edit gates via dedicated controls, link table entries to corresponding plots, and display population statistics within visualizations. In addition, flexible filtering options enable both global search and column-specific subsetting to efficiently interrogate large datasets (see footnote 1).

#### Sample Info

FlowWeb supports importing spreadsheet-based sample annotation files. A template for the spreadsheet-based sample annotation file is generated by, and can be downloaded from, the FlowWeb interface. This template contains a column with the file names extracted from the imported files and can be manually complemented with columns of sample information. Upon uploading, the software automatically searches the uploaded spreadsheet for a column whose values match the loaded rootfile identifiers. Once detected, the annotation sheet is joined to the main table by rootfile. This join strategy is particularly important because it propagates sample-level metadata not only to original files but, under the hood, also to all derived populations belonging to that sample.

As with cytometry data, sample annotation data are processed within the active session and are not stored on the server beyond the current session. Nevertheless, users are advised to ensure that uploaded metadata do not contain sensitive or identifiable information.

Beyond sample-level annotations, the software provides direct access to FCS keyword metadata. For a selected file, the full keyword list can be retrieved from the underlying flowFrame via the context-menu. The application supports both direct keyword selection and free-text search across keyword names and values, including atomic metadata fields and more complex nested objects. This enables users to inspect acquisition metadata interactively without leaving the web interface. Scientifically, this functionality is useful because FCS keywords often contain acquisition settings, instrument details, compensation-related metadata, and experiment descriptors that are relevant for quality control and tracking the origin and acquisition conditions of the data.

##### The Interactive Main Table

**Navigation:** To keep a compact display of large datasets while preserving the parent-child relationships between files and derived populations, Parent rows can be expanded or collapsed using a toggle symbol, both of a single sample subtree or of the full table (via the context menu).

**Context menus** that provide advanced options can be accessed through right-click within the table.

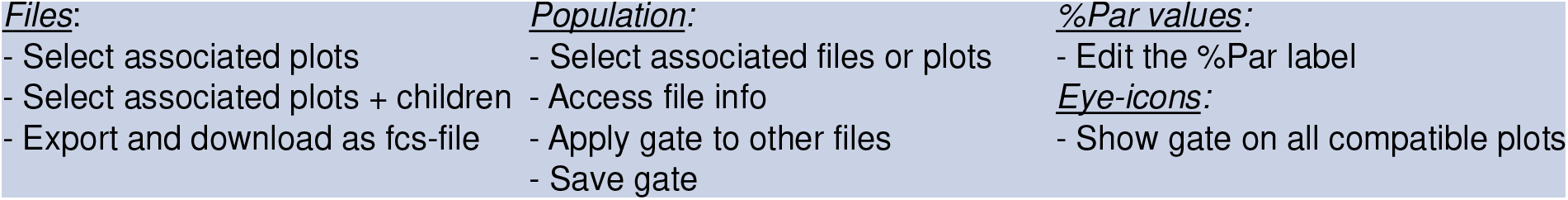

***Eye-*** and ***Pencil-*** icons adjacent to the Population name allow the visualization of the corresponding gate in the selected plots and direct access to gate editing, respectively.

**%Par** labels can be displayed in the corresponding plots by clicking the values in the table.

**Filtering** options support both global text search and column-specific filtering. Column filters can be accessed via the column header and provide either categorical selection or numeric range filtering, depending on the data type. Active filters are indicated visually in the column header.

## Functions

### Data processing

#### Quality Control

Quality control (QC) ([Data Processing] > [Quality Control] is implemented in FlowWeb using the R package flowAI[3], which provides automated detection and removal of anomalous events in flow cytometry data. The QC pipeline operates directly on flowFrame objects and is applied at the level of individual samples, while maintaining consistency with the higher-level flowSet and GatingSet structures.

QC behavior is controlled through a parameter set that includes anomaly detection modes (flow rate, signal acquisition, dynamic range), statistical thresholds, channel exclusions and output format (filtered events, QC vector, or event indices). These parameters can be manually set in the “Advanced QC parameters” modal ([Data Processing] > [Quality Control] > [Settings]) and, when logged in, can be saved for future analyses.

To improve scalability, QC is optionally applied in a chunk-wise manner, where each flowFrame is partitioned into subsets of events. This design avoids memory bottlenecks when processing large cytometry files and allows incremental reconstruction of high-quality data. These outputs are reassembled across chunks to form a cleaned dataset. The final high-quality event matrix is used to construct a new flowFrame, preserving original parameter metadata and keywords. Although chunk-wise QC improves computational scalability, its use within the flowAI framework may reduce statistical power and can therefore lead to increased event exclusion.

When QC has finished, the results appear in a new tab. The QC reports can optionally be saved and downloaded as a zip-file, which allows visual inspection of the QC results. In addition, the zip-file contains the cleaned fcs-files, which avoids repeated QC steps.

Cleaned flowFrame objects replace their corresponding entries in the flowSet and the reactive main table is updated, including the percentage of retained events.

#### Transform

Data transformation is implemented using the transformation framework provided by flowCore, which supports a wide range of mathematical transformations commonly used in flow cytometry, including:

- logarithmic
- arcsinh
- linear and scaling transforms,
- quadratic transforms
- truncation
- biexponential
- logicle

Transformations are applied at the flowSet level, ensuring that all samples within a panel-compatible group are processed consistently. Because the original (untransformed) flowSet is stored in a separate reactive registry the transformations are fully reversible, which avoids reloading files.

Because transformations modify the underlying expression matrices, all downstream analyses, including gating and visualization, operate on transformed data.

### Plot

Visualization in FlowWeb is implemented using the R package ggplot2 [4] as the primary rendering engine, combined with shiny for a dynamic user interface. Additional extensions and base statistical layers within ggplot2 are used to support a wide range of visualization modalities.

#### Individual Plots

Individual plots are generated from flowFrame objects retrieved from the internal flowSet registry. For each selected sample, the expression matrix is extracted using exprs() and converted into a standard data frame.

Each plot instance is represented by a unique identifier and associated configuration parameters, including selected channels, plot type and visualization settings, axis scales and ranges, and metadata display options.

FlowWeb supports multiple visualization modes commonly used in flow cytometry:

- Dot plots
- 1D density plots
- Density scatter plots
- Contour plots
- Box plots

All plots are constructed using layered ggplot2 grammar, allowing consistent application of themes, scales, and annotations. Hence, a wide range of the different layers of ggplot2 variables can be adjusted by the users through [Edit].

Plot instances are tracked in a centralized table, which stores layout information (position, size) and links each plot to its underlying data source. This enables multiple independent plots to coexist and be interactively positioned within the interface.

A key feature of the plotting system is its integration with the gating framework. Gate geometries are overlaid onto plots using polygon layers, ensuring that visualizations reflect the current analytical state stored in GatingSet objects. These overlays are dynamically recomputed based on the selected sample and channel configuration.

Data from the reactive metadata table allows users to display sample-level or population-level information directly within the visualization. This creates a tight coupling between data, metadata, and visual output.

#### Overlay plots

Overlay plots enable direct comparison of multiple samples within a single visualization. These plots are constructed by combining data from multiple flowFrame objects into a unified data frame, with an additional grouping variable indicating sample identity.

For each selected sample, expression values are extracted and concatenated into a single long-format data frame, allowing ggplot2 to map color, fill, or grouping aesthetics across samples.

Overlay plots support the following plot types:

- Overlay density plots
- Frequency polygons
- Scatter overlays (dot plots)
- Box plots
- Ridge plots

Overlay plots can group and color samples based on metadata columns rather than raw file identity. This is enabled by joining the combined expression data with the reactive metadata table, allowing grouping by experimental condition, population, or user-defined annotations.

#### Statistical plots

Statistical visualization is implemented as a distinct plot family that operates on aggregated tabular data rather than raw event-level measurements. These plots are generated using ggplot2 for visualization and ggpubr for statistical testing and annotation.

Statistical plots are constructed from the reactive metadata table rather than directly from flowFrame objects. For selected rows, a derived dataset is created containing sample identifiers, population labels and summary metrics such as %Par or event counts. This dataset is stored in a dedicated registry and serves as the input for all statistical visualizations. The data are reshaped into a long format suitable for ggplot2, with allows the use of sample metadata to customize the plots, such as grouping on the x-axis, fill and shape aesthetics.

Currently, two main statistical plot types are supported; bar plots, showing mean values per group with optional error bars (e.g., standard error), and stacked bar plots; hierarchical representation of populations within each sample.

#### Statistics

Statistical information can be added to Statistical plots through their context menus ([Statistics] > [Add statistics]). The “Statistics Settings” menu hosts a range of parameters that can be set by the user to tailor the statistical comparisons performed by the ggpubr package. These parameters include:

- Type of test (e.g., t-test, Wilcoxon)
- Paired vs. Unpaired design
- variance assumptions
- Multiple testing correction

When applied, the selected statistical test is automatically performed between the groups on the x-axis and results are visualized as annotated brackets above the corresponding bars.

Label formatting is configurable (e.g., stars vs. exact p-values), and non-significant results can be optionally suppressed. The system also includes logic to suggest appropriate statistical tests based on data structure (e.g., number of groups, sample size).

The “Statistics summary” modal ([Statistics] > [Show summary]) allows the user to access a summary interface that provides dataset structure (group counts), applied statistical parameters, and an overview of the underlying data. This ensures transparency and reproducibility of statistical results.

### Gating

#### Manual gating

Manual gating in FlowWeb is implemented as an interactive workflow that combines direct graphical user input with formal cytometry data structures. The core gating machinery relies on the Bioconductor packages flowCore and flowWorkspace, while Shiny provides the event-driven user interface and ggplot2 supports graphical rendering of the gating canvas. In practice, user-drawn gates are not treated as visual annotations but are converted into formal gate objects and incorporated into sample-specific gating hierarchies.

Manual gating in FlowWeb is guided by the user’s current selection (sample, channels, and gate type) and interactive input on the plot. Gates can be drawn and edited using standard geometries such as thresholds, rectangles, polygons, ellipses, and quadrants. In all cases, the drawn region is converted into a formal gate definition and added to the gating hierarchy. FlowWeb supports the following gate geometries from the flowCore package:

- 1D density gates
- Rectangle gates
- Ellipse gates
- Polygon gates
- Quadrant gates

At the data level, cytometry data are handled as *flowFrame* objects, from which expression values are accessed and gated subsets are derived. User-defined regions are translated into formal gate objects (e.g. rectangular or polygonal gates), preserving the relationship between geometric boundaries and measurement channels. These gates are embedded into a *GatingSet* / *GatingHierarchy* structure, ensuring that population relationships are maintained and consistently updated when gates are modified.

Each gated population is exported as a child population and represented both within the hierarchical gating structure and in a metadata table that records parent–child relationships, population names, event counts, and visual attributes such as gate colour. The tabular representation and the gating hierarchy are updated in parallel, ensuring consistency between views. When a gate is edited, its definition is retrieved, reconstructed in the editor, and updated in the gating hierarchy. Changes are subsequently propagated to all dependent populations, affected child populations are recomputed and their associated data and metadata are updated accordingly. This ensures that manual adjustments are consistently reflected throughout the analysis workflow.

#### Automated gating

Automated gating in FlowWeb complements manual gating by providing algorithmic methods for identifying populations based on one- or two-dimensional distributions. This part of the framework combines methods from openCyto and model-based packages such as flowClust, mixtools, and mclust. These methods are wrapped by FlowWeb so that users can select them from the interface, inspect their results in a preview panel, and then apply them to one or more samples.

FlowWeb provides automated gating methods such as:

- Cell cycle analysis
- Minimum-density thresholding
- Tail-based cutoff detection
- Quantile gating
- Two-dimensional model-based clustering

These methods are particularly appropriate for marker channels where one seeks to separate negative and positive populations, exclude debris or extreme tails, or enforce percentile-based thresholds across samples.

For two-dimensional model-based clustering, the implementation makes use of flowClust. Here, the event cloud in a selected pair of channels is modelled as a mixture of elliptical clusters. Each component is then expanded into an explicit polygonal contour corresponding to a chosen confidence level, allowing both visualization and downstream export as child populations. This approach is well-suited to cytometric populations that are approximately ellipsoidal but overlap in complex ways. FlowWeb converts these ellipsoidal models into polygonal gates for visualization and export, fitting them seamlessly into the broader gate overlay and editing infrastructure.

For the one-dimensional automated cell-cycle analysis, FlowWeb uses a hybrid strategy. The distribution is first characterized by a kernel density estimate, from which candidate peaks, intervening valleys, peak prominence, and approximate basin mass are derived. A biologically plausible G1 peak is selected from these KDE features, and a local two-component Gaussian mixture is fit in a robust window around that peak using mixtools. The fitted mixture model is used to identify the two principal DNA-content peaks and to assign G1 phase based on posterior probability. The final gates are next constructed explicitly; G1 is centered on the selected KDE peak and sized from high-confidence G1 events within its basin, whereas G2 is placed at an approximate duplication multiple of the G1 center with related width, and S phase is defined as the intervening interval between the two. Although this routine is not part of openCyto, it follows the same design principle: model a population in a statistically principled way, convert the result into explicit gates, and propagate those gates through the same export pipeline as all other methods.

## Results

To benchmark FlowWeb’s accuracy and reproducibility against established commercial platforms, we performed automated cell cycle analysis on Propidium Iodide-stained samples. To evaluate both performance and reproducibility, we analyzed samples from two conditions (Control and Treated), each with three replicates, showing clearly distinct cell cycle patterns (Figure 1). The distribution of cells over G1, S and G2 as determined by FlowWeb closely resembles the distribution as reported by FCS-Express. The results obtained by Kaluza deviated slightly, assigning fewer cells to S-phase and more to G2. Both FlowWeb as well as FCS-express demonstrated high reproducibility across the three replicates. When averaged across G1, S and G2, FlowWeb results showed a Standard Deviation of 6.69% and 3.74% in Control and Treated condition, respectively, which was comparable to that of FCS-Express (6.52% and 4.22%) (Table 1). Altogether, these results confirmed that the performance of the FlowWeb software closely matches that of commercially available software packages.

**Table 1:** Overview of the distribution of cells over G1, S and G2 phases as determined automated analysis software by cell in the cycle three packages. Note: For the Treated samples, Kaluza was only able to detect the different phases in one of three samples.

|  |  | Kaluza |  |  | FlowWeb |  |  | FCS-Express |  |  |
| --- | --- | --- | --- | --- | --- | --- | --- | --- | --- | --- |
|  |  | Mean | StDev | StDev<br>(% Mean) | Mean | StDev | StDev<br>(% Mean) | Mean | StDev | StDev<br>(% Mean) |
| Control | G1 | 71.33 | 2.36 | 3.30 | 68.86 | 2.43 | 3.53 | 67.81 | 2.55 | 3.76 |
|  | S | 1.67 | 0.47 | 28.28 | 11.71 | 1.25 | 10.66 | 16.85 | 1.43 | 8.50 |
|  | G2 | 25.00 | 2.16 | 8.64 | 16.65 | 0.98 | 5.89 | 15.34 | 1.12 | 7.31 |
|  | Average: |  |  | 13.41 |  |  | 6.69 |  |  | 6.52 |
| Treated | G1 | 24.00 | N/A | N/A | 26.69 | 0.98 | 3.68 | 15.18 | 1.32 | 8.72 |
|  | S | 1.00 | N/A | N/A | 23.00 | 1.22 | 5.32 | 38.03 | 0.89 | 2.33 |
|  | G2 | 74.00 | N/A | N/A | 46.55 | 1.03 | 2.20 | 46.78 | 0.75 | 1.61 |
|  | Average: |  |  | N/A |  |  | 3.74 |  |  | 4.22 |

**Figure 1:**
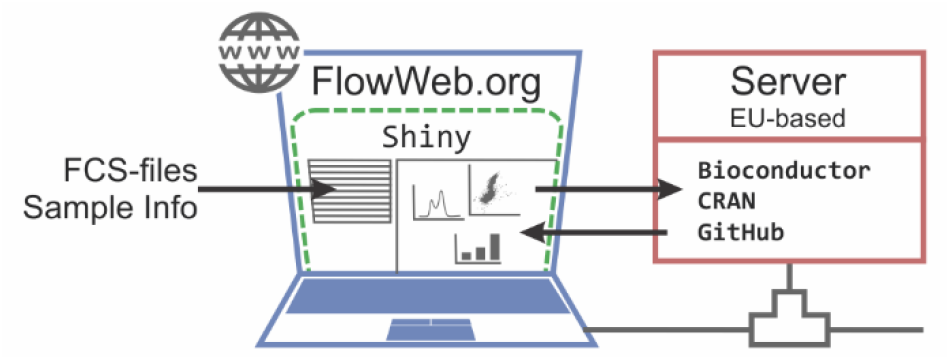
**Frontend–Backend Architecture Overview:** browser-based interface with server-side analysis using open-source R packages from CRAN, Bioconductor and GitHub.

**Figure 2:**
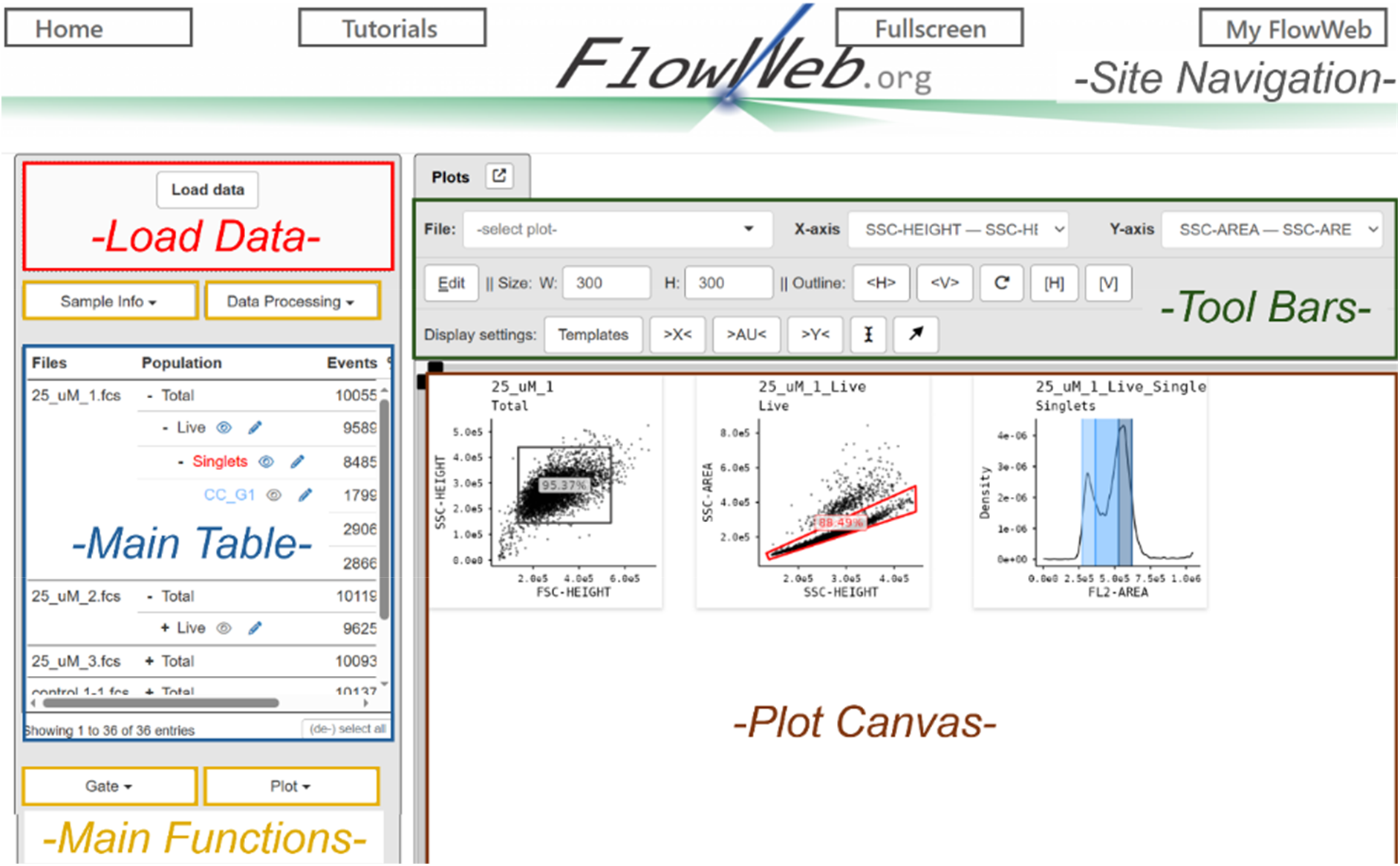
**Overview of the FlowWeb interface**, showing the main reactive components of the web-based analysis environment, including the main table, plot canvas, and associated control panels.

**Figure 3.**
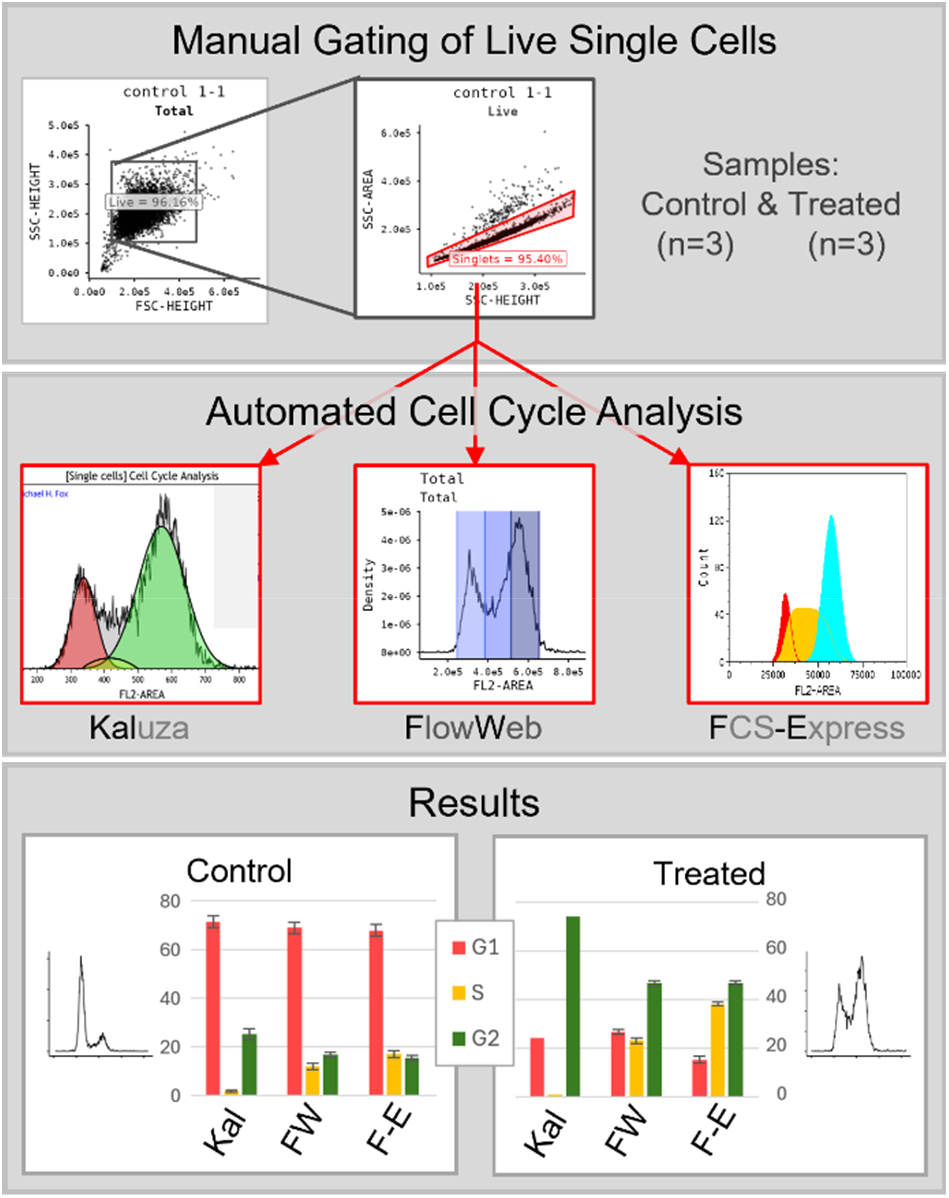
A dataset of 6 samples (3 x Control, 3 x Treated) was first manually cleaned in FlowWeb by excluding debris and doublets. The percentage of cells in the three phases of the cell cycle was then quantified using automated cell cycle analysis pipelines from three different software packages. Representative DNA content profiles are shown along side the graphs. Error bars represent the standard deviation of the mean.

As in the manual case, flowCore provides the cytometry containers and gate classes, flowWorkspace manages hierarchical integration and recomputation, shiny orchestrates the interactive workflow, and ggplot2 renders previews and final overlays.

## My FlowWeb

Flow Web provides a user-specific environment (“My FlowWeb”) that allows storage and reuse of analysis configurations across sessions. This functionality is implemented through a hybrid architecture combining a reactive R-based frontend through shiny, with a backend layer implemented as a custom WordPress plugin.

The system enables users to store and retrieve three distinct classes of objects: (i) quality control (QC) presets, (ii) plot templates, and (iii) gating definitions. All objects are associated with authenticated user accounts and are versioned and timestamped in a relational database. Communication between the Shiny application and the backend is performed using the httr2 HTTP client, with authentication enforced through WordPress session cookies and nonce-based request validation. The different objects are stored in a dedicated relational table within WordPress, with each record linked to a specific user ID. This design allows different aspects of the workflow to be treated as reusable configuration objects, decoupled from specific datasets while remaining fully reproducible.

**QC presets**; a list of QC parameter configurations used during preprocessing and quality control steps, allowing users to standardize workflows across datasets.

**Plot templates**; includes amongst others plot type and rendering parameters (e.g., density adjustment, binning), aesthetics (colors, transparency, contour properties), axis configuration (scales, labels, visibility),annotation settings (header and subheader formatting), and legend and display options. Plot templates are portable across datasets and allow rapid replication of complex visualization settings without reconfiguration.

**Gates**; encode explicit geometric and analytical gating objects that can be reapplied across datasets. These objects integrate tightly with the cytometry infrastructure provided by flowCore and flowWorkspace. Within the app, logged-in users can retrieve an overview of saved gates from the reactive storage on demand. This overview includes information related to gate type, associated channels and optional metadata (color, source file, versioning, edit history).

User-specific objects stored within “My FlowWeb,” including plot templates, QC presets, and gating definitions, consist of analytical parameters such as channel selections, visualization settings, thresholds, and gate geometries. These objects do not store raw cytometry data and are not intended to contain patient-identifiable information. However, under frameworks such as the GDPR, information may be considered personal if it can directly or indirectly identify an individual. Users are therefore responsible for ensuring that no sensitive information is embedded in user-defined labels, annotations, or metadata associated with these stored configurations.

